# Vangl2 acts in distinct cell types to establish bidirectional hair-bundle polarity and maintains tissue-wide alignment in zebrafish neuromasts

**DOI:** 10.64898/2026.08.24.746764

**Authors:** Swarna Jeewajee, Francesco Gianoli, Maria Jussila, Brian Ciruna, Aaron Steiner, Adrian Jacobo, A. J. Hudspeth

## Abstract

The conserved core planar cell polarity (PCP) pathway orients cells and subcellular structures within an epithelium through asymmetric protein localization and intercellular communication(1–4). In vestibular organs and lateral-line neuromasts, mechanosensory hair cells are interspersed among support cells and form opposing hair-bundle orientations along a shared axis, enabling bidirectional sensitivity to head motion and water flow, respectively(5–9). In zebrafish neuromasts, Notch-mediated lateral inhibition gives rise to two hair-cell populations, distinguished by differential Emx2 expression, that orient their cell-intrinsic polarity machinery differently relative to a PCP-dependent tissue-wide axis(10–14). However, it remains unclear how PCP proteins are organized across hair cells and support cells to achieve both opposing hair-bundle orientations and tissue-wide alignment, and whether PCP signaling remains required after hair-bundle polarity is established. Combining quantitative spatial mapping of the core PCP protein Vangl2 with cell-type-specific and temporally controlled protein degradation, we show that hair cells and support cells make distinct yet coordinated contributions to the polarized Vangl2 organization within neuromasts and to bidirectional hair-bundle polarity. Support-cell Vangl2 facilitates tissue-wide alignment of hair bundles along the anteroposterior axis, whereas hair-cell Vangl2 is required to generate opposing hair-bundle orientations along this axis. Vangl2 degradation after hair bundles have formed disrupts their tissue-wide alignment, showing that planar polarity is actively maintained rather than fixed after establishment. Together, these findings reveal how Vangl2-dependent PCP signaling is distributed across distinct cell types within a heterogeneous epithelium to generate opposing polarity outcomes and remains necessary to preserve tissue-level planar organization.

## Results

### Both support cells and hair cells contribute to polarized Vangl2 organization in neuromasts

The core planar cell polarity (PCP) pathway coordinates the orientation of cells and subcellular structures within epithelia to ensure proper tissue architecture and function(2, 3). Core PCP proteins, including Frizzled, Dishevelled, Prickle, Van Gogh/Vangl, and Celsr/Flamingo, become asymmetrically distributed within cells and interact across cell-cell junctions to propagate directional information between neighbors(1, 4, 15). Much of our understanding of PCP signaling has emerged from epithelia composed of equivalent cells that adopt uniform polarity. However, bidirectionally sensitive mechanosensory epithelia, including vestibular organs of the inner ear and zebrafish lateral-line neuromasts, pose two distinct organizational challenges: hair cells are interspersed with support cells, and their mechanosensory hair bundles must adopt opposing orientations while remaining aligned along a common polarity axis to detect stimuli from opposite directions for balance and rheotaxis, respectively(6–9) (Fig. 1A).

**Fig. 1.**
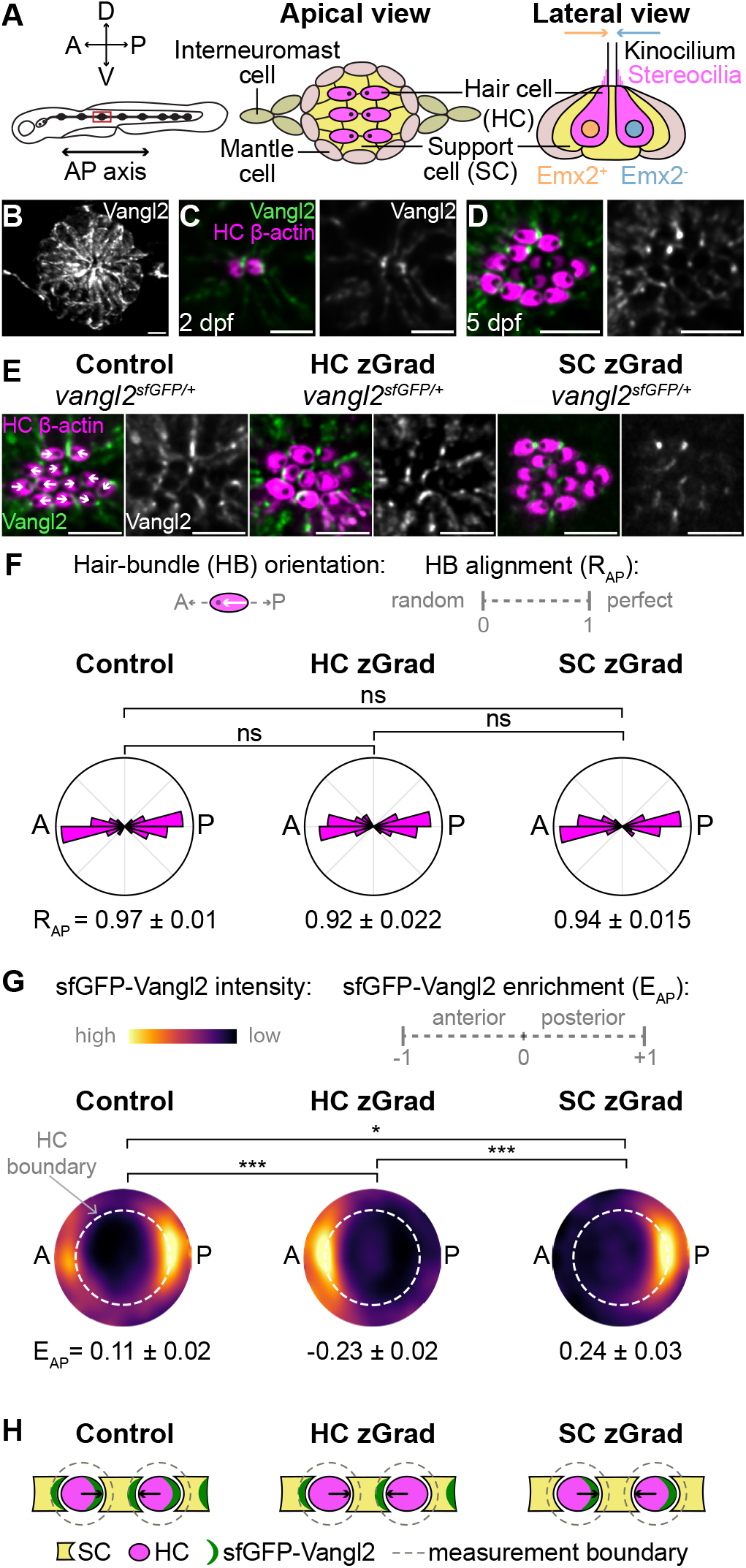
Both support cells and hair cells contribute to polarized Vangl2 organization in neuromasts. (A) Schematic of the posterior lateral line indicating the anteroposterior (AP) and dorsoventral axes and the cellular organization of a neuromast (red box). A lateral-view schematic illustrates hair cells (HCs) with oppositely oriented hair bundles (HBs), surrounded by support cells (SCs). The oppositely oriented HBs detect water flow in opposite directions and correspond to differential Emx2 expression in HC nuclei. (B) Representative maximum-intensity projection of a *vangl2*^sfGFP/+^ neuromast at 3 days post fertilization (dpf), illustrating broad Vangl2 expression. (C, D) Representative apical images of *vangl2*^sfGFP/sfGFP^; *Tg(myo6b:actb1-mScarletI)* neuromasts at 2 dpf (C) and 5 dpf (D), illustrating sfGFP-Vangl2 organization around HCs at early and later stages, respectively. HC *β*-actin marks actin-rich hair bundles (in magenta) but not the kinocilium, and sfGFP-Vangl2 is shown in green or grayscale. (E) Representative apical images of 5 dpf *vangl2*^sfGFP/+^; *Tg(myo6b:actb1-mScarletI)* neuromasts under control conditions or after zGrad-mediated degradation of sfGFP-Vangl2 in HCs (HC zGrad, *Tg(myo6b:zGrad-2A-mCherry)*) or SCs (SC zGrad, *Tg(she:zGrad-2A-mCherry)*). White arrows illustrate HB orientations. (F) Polar histograms of HB orientations in 5 dpf neuromasts. AP alignment was quantified per neuromast using axial resultant length, *R*_AP_ (0 = random, 1 = perfectly aligned). (G) Shape-normalized sfGFP-Vangl2 maps showing relative signal around HC apical boundaries in 5 dpf neuromasts. Color shows the spatial distribution of Vangl2 enrichment (brighter color means higher enrichment) relative to the neuromasts’ average; all panels share one scale. Dashed circles indicate the normalized HC boundary. *E*_AP_ quantifies AP sfGFP-Vangl2 enrichment; positive values indicate posterior enrichment, and negative values anterior enrichment. (H) Schematic of Vangl2 distributions after cell-type-specific degradation. For F and G, n = 113 HCs/13 neuromasts/4 fish for control, 136 HCs/15 neuromasts/4 fish for HC zGrad, and 132 HCs/11 neuromasts/7 fish for SC zGrad. Data are mean ± SEM across neuromasts. Statistical significance was assessed using Kruskal– Wallis tests followed by pairwise Mann–Whitney U tests with Holm correction on neuromast-level *R*_AP_ or *E*_AP_ values. For *R*_AP_, Holm-corrected pairwise tests: control vs HC zGrad, P = 0.136; control vs SC zGrad, P = 0.136; HC zGrad vs SC zGrad, P = 0.977. For *E*_AP_, Holm-corrected pairwise tests: control vs HC zGrad, P = 5.52 × 10^−5^; control vs SC zGrad, P = 1.14 × 10^−2^; HC zGrad vs SC zGrad, P = 4.08 × 10^−5^. ns, not significant; *P < 0.05, ***P < 0.001. Scale bars, 5 µm. See also Figure S1.

Previous results have shown that hair bundle orientation is disrupted in vangl2 mutant fish, or when Vangl2 is over-expressed in hair cells(12). Here, we define how Vangl2 is organized across hair cells and support cells in zebrafish neuromasts (Fig. 1A) by visualizing endogenously tagged sfGFP-Vangl2(16). sfGFP-Vangl2 was broadly enriched within neuromasts (Fig. 1B) and localized asymmetrically around the apical perimeters of hair cells along the anteroposterior (AP) axis at both early and later stages of maturation (Fig. 1B–D). To determine whether polarized Vangl2 organization occurs in hair cells, support cells, or both, we induced sfGFP-Vangl2 degradation in each cell type using cell-type-specific expression of the degron zGrad(17) in *vangl2*^sfGFP/+^ fish (Fig. 1E), which was confirmed by complementary zGrad-2A-mCherry expression and residual sfGFP-Vangl2 distributions in distinct cellular compartments (Fig. S1A–C). Hair-bundle orientations in the *vangl2*^sfGFP/+^ neuromasts were not detectably different from controls (Fig. 1F), allowing us to assess cell-type-specific contributions to polarized Vangl2 localization in a genetic background where PCP signaling remained broadly functional. Because hair cells vary in size and shape, we fitted an ellipse to each traced apical outline, then rescaled the cell and the fluorescence around to a unit circle (Fig. S1B). Interior, boundary, and exterior regions then occupy the same radial coordinates in every cell, with the boundary at *r* = 1, so that Vangl2 enrichment inside and outside hair cells can be averaged and compared across cells. These shape-normalized maps revealed the relative enrichment of sfGFP-Vangl2 around hair-cell boundaries, and the directional bias of this distribution was quantified using the anteroposterior enrichment index, *E*_AP_, for which negative and positive values indicate anterior and posterior bias, respectively (see Methods). Under control conditions, Vangl2 was enriched at both poles along the anteroposterior axis (Fig. 1G). Degradation in either hair cells or support cells reduced the overall Vangl2 signal but left complementary polarized distributions: hair-cell-specific degradation left an anterior-biased Vangl2 distribution attributable to support cells, whereas support-cell-specific degradation left a posterior-biased distribution attributable to hair cells (Fig. 1G). Together, these results indicate that both hair cells and surrounding support cells contribute to the polarized distribution of Vangl2 in neuromasts (Fig. 1H).

### Support-cell Vangl2 facilitates tissue-wide alignment but is insufficient to orient hair bundles in the absence of hair-cell Vangl2

To assess the functional implications of these cell-type-specific Vangl2 contributions, we performed zGrad-mediated degradation in *vangl2*^sfGFP/sfGFP^ larvae, in which both Vangl2 alleles are sfGFP-tagged and therefore susceptible to degradation. Under control conditions, Vangl2 was enriched at both poles along the anteroposterior axis (Fig. 2A, B), and hair bundles were tightly aligned with this axis (Fig. 2C). Strikingly, cell-type-specific Vangl2 degradation produced distinct effects on Vangl2 localization and hair-bundle polarity. Support-cell-specific degradation reduced the directional bias in the residual hair-cell Vangl2 distribution (Fig. 2A, B) and broadly disrupted hair-bundle alignment (Fig. 2C). By contrast, despite the persistence of an anterior-biased Vangl2 distribution in the surrounding support cells (Fig. 2A, B), hair-cell-specific degradation disrupted the alignment of hair bundles and produced an anterior orientation bias (Fig. 2C). Thus, support-cell Vangl2 facilitates tissue-wide hair-bundle alignment but is insufficient to properly orient hair bundles in the absence of hair-cell Vangl2.

**Fig. 2.**
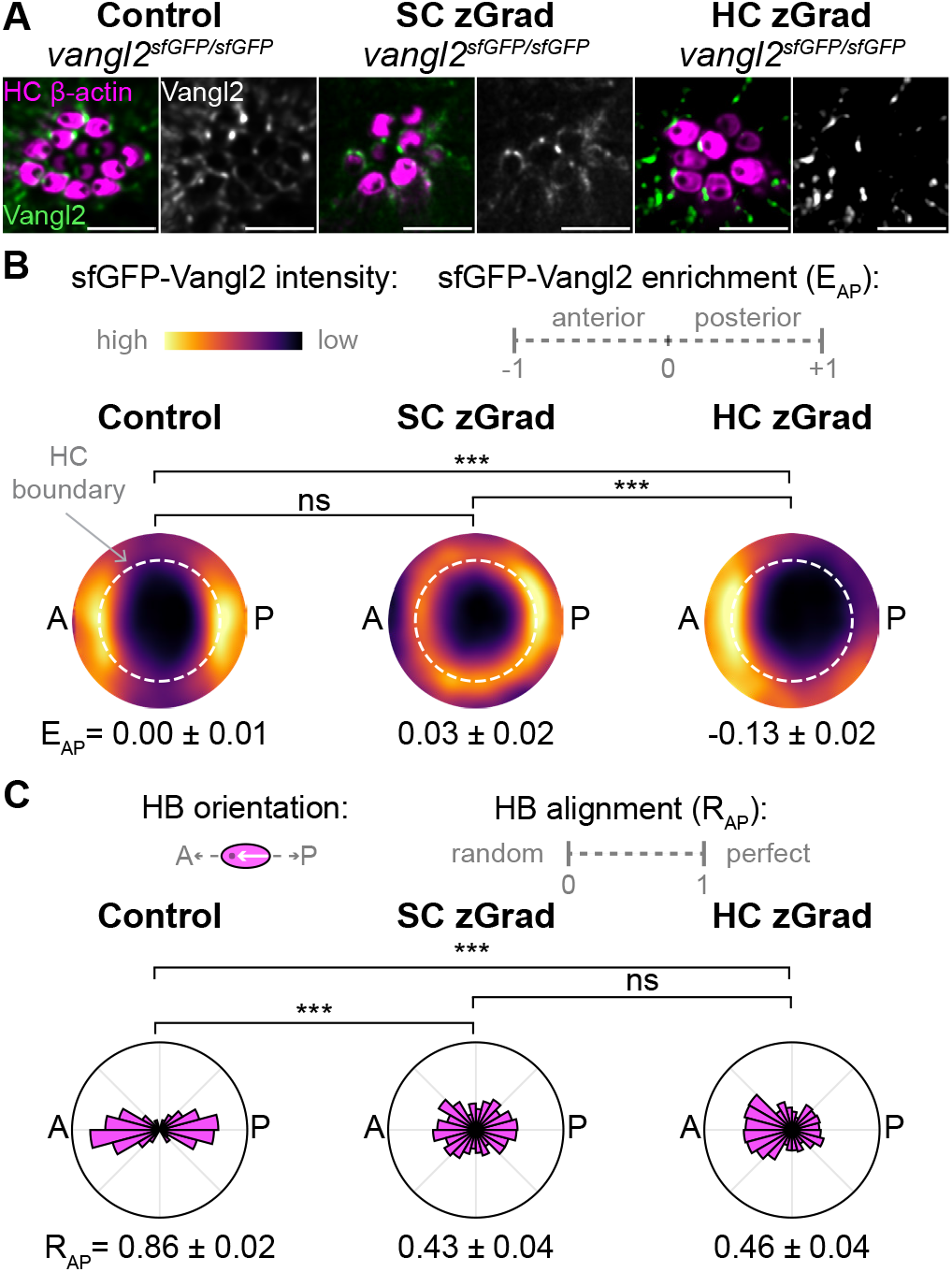
Cell-type-specific Vangl2 degradation has distinct effects on Vangl2 organization and bidirectional hair-bundle polarity. (A) Representative apical images of 5 dpf *vangl2*^sfGFP/sfGFP^; *Tg(myo6b:actb1-mScarletI)* neuromasts under control conditions or with zGrad-mediated degradation of sfGFP-Vangl2 in hair cells (HCs; HC zGrad, *Tg(myo6b:zGrad-2A-mCherry)*) or support cells (SCs; SC zGrad, *Tg(she:zGrad-2A-mCherry)*). Merged images show HC *β*-actin (magenta) and sfGFP-Vangl2 (green); adjacent grayscale images show sfGFP-Vangl2 alone. (B) Shape-normalized sfGFP-Vangl2 maps showing relative signal around HC apical boundaries in 5 dpf neuromasts. Color shows the spatial distribution of Vangl2 enrichment (brighter color means higher enrichment) relative to the neuromasts’ average; all panels share one scale. Dashed circles indicate the normalized HC boundary. *E*_AP_ quantifies AP sfGFP-Vangl2 enrichment; positive values indicate posterior enrichment, and negative values anterior enrichment. (C) Polar histograms of hair-bundle (HB) orientations in 5 dpf neuromasts. AP alignment was quantified per neuromast using axial resultant length, *R*_AP_ (0 = random, 1 = perfectly aligned). For B and C, n = 234 HCs/22 neuromasts/7 fish for control, 276 HCs/35 neuromasts/10 fish for HC zGrad, and 229 HCs/21 neuromasts/6 fish for SC zGrad. Data are mean ± SEM across neuromasts. Statistical significance was assessed using Kruskal–Wallis tests followed by pairwise Mann–Whitney U tests with Holm correction using neuromast-level *E*_AP_ or *R*_AP_ values. For *E*_AP_, Holm-corrected pairwise tests: control vs HC zGrad, P = 1.80 × 10^−6^; control vs SC zGrad, P = 0.179; HC zGrad vs SC zGrad, P = 1.8 × 10^−6^. For *R*_AP_, Holm-corrected pairwise tests: control vs HC zGrad, P = 5.69 × 10^−8^; control vs SC zGrad, P = 2.51 × 10^−7^; HC zGrad vs SC zGrad, P = 0.977. ns, not significant; ***P < 0.001. Scale bars, 5 µm. See also Figure S1.

### Hair-cell Vangl2 is required to establish opposing hair-bundle orientations

In neuromasts, Notch-mediated lateral inhibition generates two hair-cell populations distinguished by the differential expression of the transcription factor Emx2(10, 11, 14, 18). These populations orient their cell-intrinsic polarity machinery in opposite directions relative to a Vangl2-dependent tissue axis(12, 13, 19, 20): in wild-type neuromasts, Emx2-negative hair cells orient their bundles anteriorly, whereas Emx2-positive hair cells orient them posteriorly. We therefore hypothesized that the anterior bias following hair-cell-specific Vangl2 degradation reflected a differential effect on these two populations. To test this hypothesis, we used immunohistochemistry to identify Emx2-negative and Emx2-positive hair cells (Fig. S2) and analyzed hair-bundle orientations within each population. Under control conditions, Emx2-positive and Emx2-negative hair cells adopted opposing hair-bundle orientations aligned with the anteroposterior axis (Fig. 3A). Following hair-cell-specific Vangl2 degradation, hair-bundle orientations in Emx2-negative hair cells remained largely confined to their expected anterior quadrant, whereas Emx2-positive hair cells displayed a broader distribution that spanned both anterior and posterior quadrants and showed reduced anteroposterior alignment (Fig. 3B). By contrast, Vangl2 degradation within support cells affected hair-bundle orientation more uniformly across both populations, consistent with a broader tissue-level perturbation rather than a cell type-specific defect (Fig. 3C). Thus, support-cell Vangl2 facilitates anteroposterior alignment of hair bundles across both hair-cell populations, whereas hair-cell-intrinsic Vangl2 is required to establish opposing hair-bundle orientations (Fig. 3D).

**Fig. 3.**
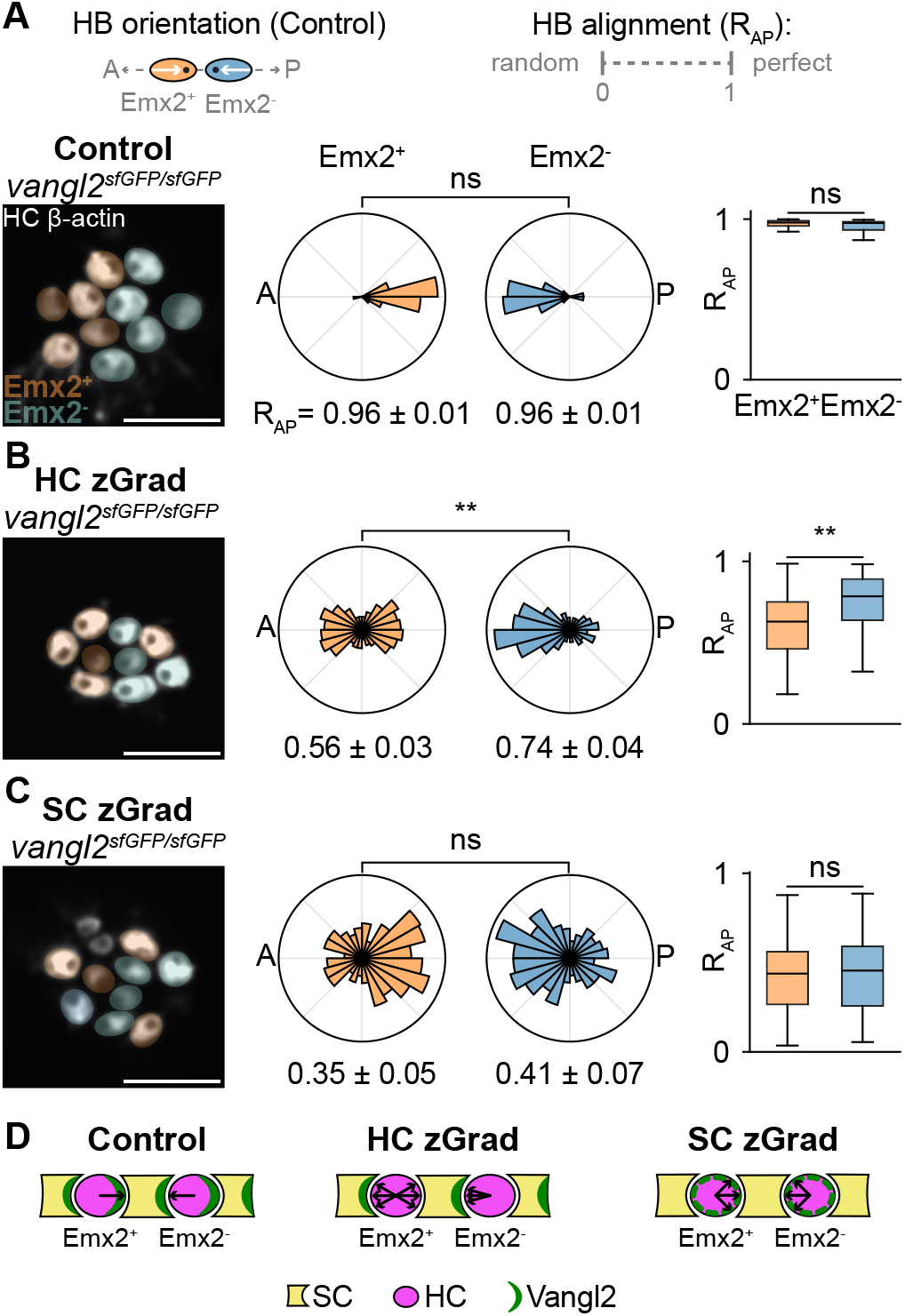
Hair-cell-specific Vangl2 degradation differentially affects hair-bundle polarity in Emx2-positive and Emx2-negative hair cells. (A–C) Emx2-specific hair-bundle polarity in 5 dpf *vangl2*^sfGFP/sfGFP^; *Tg(myo6b:actb1-mScarletI)* neuromasts under control conditions (A), after zGrad-mediated degradation of sfGFP-Vangl2 in hair cells (HCs; B, HC zGrad, *Tg(myo6b:zGrad-2A-mCherry)*), or after degradation in support cells (SCs; C, SC zGrad, *Tg(she:zGrad-2A-mCherry)*). The top schematic in A indicates the expected opposing hair-bundle (HB) orientations of Emx2^+^ and Emx2^−^ HCs in control conditions. Left panels show representative apical images with HCs color-coded by Emx2 expression: Emx2^+^ HCs, orange; Emx2^−^ HCs, blue. Colors indicate Emx2 identity, independent of measured HB orientation. Middle panels show polar histograms of HB orientations for Emx2^+^ and Emx2^−^ HCs. Right panels show paired neuromast-level AP alignment, quantified separately for Emx2^+^ and Emx2^−^ HCs using axial resultant length, *R*_AP_ (0 = random, 1 = perfectly aligned). (D) Schematic summarizing Vangl2 localization and HB orientation phenotypes under each condition. Arrows indicate predominant HB orientations. For A–C, n = 315 HCs/33 neuromasts/16 fish for control, 555 HCs/58 neuromasts/32 fish for HC zGrad, and 282 HCs/23 neuromasts/13 fish for SC zGrad. Data are mean ± SEM across neuromasts. Statistical significance was assessed using two-tailed Wilcoxon signed-rank tests comparing paired neuromast-level *R*_AP_ values for Emx2^+^ and Emx2^−^ HCs within each condition. Exact p-values: control, P = 0.405; HC zGrad, P = 0.00308; SC zGrad, P = 0.823. ns, not significant; **P < 0.01. Scale bars, 5 µm. See also Figure S2.

### Vangl2 is required beyond polarity establishment to maintain hair-bundle alignment

Having established the distinct cell-type-specific requirements for Vangl2 in generating bidirectional hair-bundle polarity, we next asked whether Vangl2 is required only during initial polarization or remains necessary after hair-bundle polarity has been established. To distinguish these temporal requirements, we used heat-shock-induced activation of zGrad to degrade Vangl2 either at 1 day post fertilization (dpf), before hair-bundle polarity is established, or at 5 dpf, after hair bundles have become aligned along the anteroposterior axis (Fig. 4A). Global Vangl2 degradation at 1 dpf disrupted hair-bundle orientation when assessed at 5 dpf (Fig. 4B, C). Surprisingly, global Vangl2 degradation at 5 dpf also disrupted hair-bundle orientation by 7 dpf (Fig. 4B, C).

**Fig. 4.**
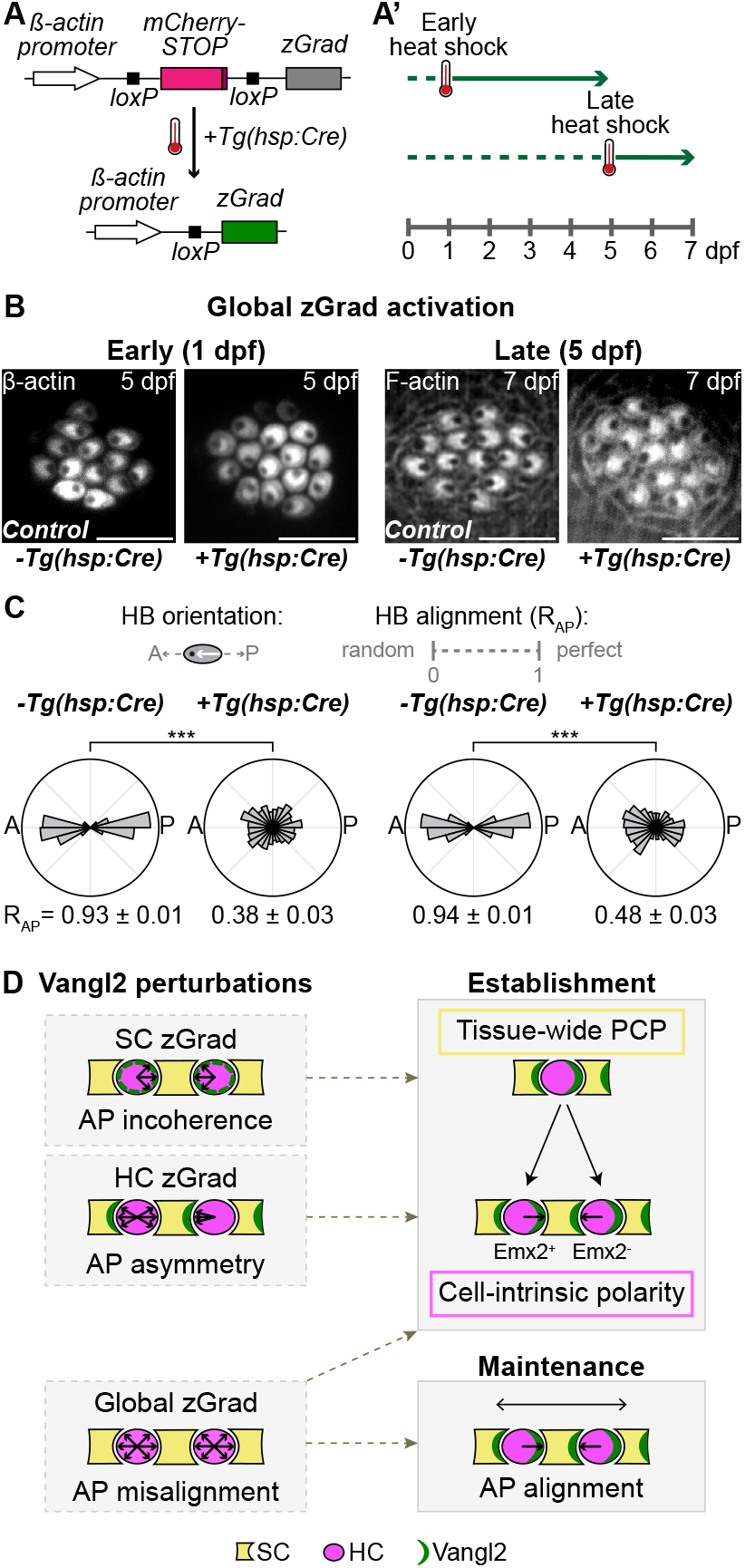
Vangl2 is required not only to establish but also to maintain hairbundle polarity. (A) Schematic of heat shock-induced global zGrad activation. In *Tg(βactin2:loxP-mCherry-STOP-loxP-zGrad)* larvae, zGrad expression is blocked by an intervening mCherry-STOP cassette. Heat shock-induced Cre expression from *Tg(hsp:Cre)* excises the STOP cassette through recombination at loxP sites, placing zGrad under the control of the *β*-actin promoter and halting mCherry expression. (A^*′*^) Experimental timeline. Heat shock was applied at 1 dpf to perturb Vangl2-dependent signaling during hair-bundle (HB) polarity establishment, or at 5 dpf to test whether Vangl2 is required to maintain established polarity. Arrowheads indicate imaging time points. (B) Representative apical images of *vangl2*^sfGFP/sfGFP^; *Tg(βactin2:loxP-mCherry-STOP-loxP-zGrad)* neuromasts with or without *Tg(hsp:Cre)* following early or late heat-shock. Early heat-shocked larvae were imaged at 5 dpf, and late heat-shocked larvae were imaged at 7 dpf. HBs were visualized using HC *β*-actin in *Tg(myo6b:actb1-mScarletI)* larvae or F-actin labeling with phalloidin. (C) Polar histograms of HB orientations. AP alignment was quantified per neuromast using axial resultant length, *R*_AP_ (0 = random, 1 = perfectly aligned). For C, n = 211 HBs/15 neuromasts/6 fish for early *Tg(hsp:Cre)*, 447 HBs/44 neuromasts/18 fish for early +*Tg(hsp:Cre)*, 226 HBs/16 neuromasts/9 fish for late −*Tg(hsp:Cre)*, and 179 HBs/14 neuromasts/7 fish for late +*Tg(hsp:Cre)*. Data are mean ± SEM across neuromasts. Statistical significance was assessed using Kruskal–Wallis tests followed by pairwise Mann–Whitney U tests with Holm correction, using neuromast-level *R*_AP_ values. Holm-corrected pairwise tests: early −*Tg(hsp:Cre)* vs early +*Tg(hsp:Cre)*, P = 4.85 × 10^−8^; late −*Tg(hsp:Cre)* vs late +*Tg(hsp:Cre)*, P = 1.43 × 10^−5^. ***P < 0.001. (D) Model summarizing Vangl2-dependent organization of neuromast planar polarity. SC Vangl2 promotes tissue-wide AP alignment, HC Vangl2 enables Emx2^+^ and Emx2^−^ HCs to adopt opposing HB orientations, and Vangl2-dependent signaling maintains AP alignment after polarity is established. Scale bars, 5 µm. See also Figure S3.

Because neuromasts continue to add hair cells between 5 and 7 dpf(21, 22), we verified whether hair cells present at the time of late Vangl2 degradation persisted to 7 dpf. Larvae were fluorescently labeled immediately after heat shock at 5 dpf using FM1-43, a styryl dye that enters hair cells through their mechanotransduction channels (Fig. S3A). Neuromasts were then imaged either 1 h or 48 h later to visualize initial dye incorporation or dye retention, respectively (Fig. S3B, C). Hair cells that retained the dye by 7 dpf displayed misoriented hair bundles, consistent with previously polarized hair bundles subsequently becoming misaligned with the anteroposterior axis (Fig. S3C). Together, these results demonstrate that Vangl2 is required not only to establish hair-bundle polarity but also to maintain tissue-wide alignment (Fig. 4D).

## Discussion

Bidirectionally sensitive mechanosensory epithelia must generate opposing hair-bundle orientations, maintain their alignment along a common tissue axis, and preserve this organization after it has been established(5–7, 9). Our findings show that, in zebrafish neuromasts, these challenges are met through distinct yet coordinated functions of the core PCP protein Vangl2 across intercalated hair cells and support cells, together with its continued activity after polarity establishment. Support-cell Vangl2 facilitates tissue-wide anteroposterior alignment, whereas hair-cell-intrinsic Vangl2 enables the two hair-cell populations to adopt opposing orientations relative to this axis (Figs. 2 and 3). Temporally controlled degradation further reveals that Vangl2 remains required after polarity has been established to preserve tissue-wide alignment (Fig. 4).

Cell-type-specific Vangl2 degradation revealed that distinct cell types can coordinate to generate a shared polarity pattern while serving different functions in tissue polarization (Figs. 2 and 3). Because support cells surround and outnumber hair cells, their collective Vangl2 distribution is well positioned to confer an anteroposterior polarity axis across the neuromast(13); accordingly, support-cell-specific degradation disrupted the polarized organization of Vangl2 within hair cells and broadly misaligned hair bundles. By contrast, after hair-cell-specific degradation, support-cell Vangl2 distribution around hair cells remained organized along the anteroposterior axis, yet opposing hair-bundle orientations were disrupted. This uncoupling suggests that the tissue-wide Vangl2 pattern facilitated by support cells is insufficient on its own to generate bidirectional hair-bundle polarity and must be coupled to Vangl2 function within hair cells.

The preferential disruption of Emx2-positive hair cells provides further insight into the hair-cell-specific requirement for Vangl2 in establishing opposing hair-bundle orientations (Fig. 3). Previous work has shown that STK32A contributes to hair-bundle orientation in Emx2-negative hair cells, whereas Emx2-dependent GPR156–G*α*i signaling reverses this orientation in Emx2-positive hair cells(14, 19, 20, 23–27). Because GPR156 has been reported to associate with VANGL proteins(27), this reversal mechanism may rely more strongly on hair-cell-intrinsic Vangl2. By contrast, the relative preservation of hair-bundle orientation in Emx2-negative hair cells may reflect the ability of STK32A-associated machinery within these cells to respond to PCP information retained in the surrounding support cells. Determining how these identity-specific polarity programs interact with core PCP components in each hair-cell population will be an important direction for future work.

Temporally controlled degradation of Vangl2 after hair bundles had acquired coordinated orientation disrupted tissue-wide alignment, indicating that hair-bundle polarity is not passively retained but remains dependent on Vangl2 after its initial establishment (Fig. 4). Previous studies have followed later outcomes of Vangl2 disruption initiated before proper polarity establishment, including the progressive reorientation of initially misoriented cochlear hair cells(27, 28). However, to our knowledge, no previous study has involved the disruption of Vangl2 or another core PCP component after hair-bundle polarity had been properly established. In neuromasts, this continued requirement may reflect the need to preserve polarized interactions between hair cells and support cells as the organ grows, remodels, and incorporates newly differentiating hair cells(21, 22).

Together, our findings extend classical PCP models to a heterogeneous epithelium by showing how a core PCP component acts across distinct cell types to generate opposing polarity outcomes and remains necessary for tissue-wide planar organization.

## Methods

### Experimental model and subject details

Zebrafish work was performed in accordance with animal protocol 22081-H reviewed and approved by The Rockefeller University’s Institutional Animal Care and Use Committee. Adult zebrafish (*Danio rerio*) were maintained under standard conditions. Naturally spawned and fertilized eggs were maintained in system water treated with 0.5 mg/L methylene blue (blue water) at 28.5 °C on a 14-hour light/10-hour dark cycle. All experiments were performed on 1-7 dpf larvae, before sex can be determined. Sample sizes are reported in the corresponding figure legends.

Wild-type TU zebrafish were obtained from the Zebrafish International Resource Center. The following published zebrafish lines were used: *vangl2*^sfGFP^, *Tg(βactin2:loxP-mCherry-STOP-loxP-zGrad)*(16), and *Tg(hsp:Cre)* (29). *Tg(myo6b:actb1-mScarletI)* was generated by replacing the eGFP open reading frame in *Tg(myo6b:actb1-EGFP)*(19) with mScarletI (GenScript Biotech Corporation). The transgenic lines *Tg(myo6b:zGrad-2A-mCherry)* and *Tg(she:zGrad-2A-mCherry)* were generated using the Gateway-based Tol2 kit by combining either p5’E-myo6b (30) or p5’E-she (31), pME-zGrad, p3’E-2A-mCherry (Tol2kit), and pDestTol2pA2 (Tol2kit) entry plasmids (32). To generate the pME-zGrad, the zGrad open reading frame was amplified from pCS2(+)-zGrad (a kind gift from H. Knaut; Addgene #119716) to generate flanking attB1 and attB2 recombination sites using the following primers: attb1zGrad F 5’-GGGGACAAGTTTGTACAAA-AAAGCAGGCTATGGAGACGGAGATGGAG-3’ and zGradattb2 R 5’-GGGGACCACTTTGTACAAGAAAGCTGGGT-AGCTGGAGACGGTGACCTG-3’. The amplicon was gel extracted and cloned into the pDONR221 vector. All DNA plasmids were verified through long-read sequencing (Plasmidsaurus) and 2.0 nL of each plasmid was injected at a concentration of 2530 ng/µL into single-cell embryos along with 125 ng/µL of Tol2 transposase mRNA synthesized in vitro in the laboratory. Injected embryos were raised until sexual maturity, and adult founders with germline transmission of the transgene were identified to create a stable line.

#### Live imaging

Live larvae were anesthetized in 500 µM tricaine (MS-222; Syndel), embedded in 1% low–melting-point agarose in glass-bottom dishes, and overlaid with anesthetic solution. Images were acquired on a microlens-based super-resolution confocal microscope (VT-iSIM, VisiTech International) using a 100× silicone-oil objective. Z-stacks were collected at 300-nm intervals and deconvolved using classic maximum likelihood estimation (MLE) in Huygens software.

#### Immunofluorescence microscopy

Whole-mount larvae (5 dpf) were fixed overnight at 4 °C in 4% formaldehyde in phosphate-buffered saline (PBS) and washed four times in PBS containing 0.05% Tween 20 (PBST). Larvae were permeabilized in prechilled acetone at −20 °C for 5 minutes, washed three times in 0.05% PBST for 5 minutes each, and blocked for 2 h at room temperature with gentle nutation in 1.5% bovine serum albumin and 1.5% goat serum in 0.05% PBST. Samples were incubated overnight at 4 °C with rabbit anti-Emx2 antibody (1:200), washed four times in 0.05% PBST for 15 minutes each, and incubated overnight with Alexa Fluor 633– conjugated goat anti-rabbit secondary antibody (1:250). Larvae were washed four times in 0.05% PBST for 15 minutes each and stored at 4 °C in VectaShield (Vector Laboratories).

Fixed larvae were mounted on glass slides and imaged using a microlens-based super-resolution confocal microscope (VT-iSIM, VisiTech International) with a 100× silicone-oil objective. Z-stacks were collected at 300-nm intervals and deconvolved using classic MLE in Huygens software.

#### Heat shock treatments

Embryos were heat shocked at the indicated time points by incubation in a 37 °C water bath for 1 h, then returned to 28.5 °C until subsequent imaging at 5 or 7 dpf.

#### FM 1-43 dye treatment

Larvae were treated with 3 µM FM 1-43 dye for 45 seconds and washed three times for 1 minute each, then returned to 28.5 °C under dark conditions until subsequent imaging at 7 dpf.

### Quantification and statistical analysis

All analyses were performed using ImageJ and custom Python scripts. Statistical tests, sample sizes, and significance values are reported in the corresponding figure legends and described below. Across all analyses, the neuromast was treated as the independent unit of replication, and the number of contributing fish is reported alongside n in each figure legend. Random allocation and stratification were not applicable because experimental groups were defined by genotype and transgene combination. Sample sizes were not predetermined using statistical methods. Hair cells with ambiguous hair-bundle morphology or orientation were excluded from analyses.

#### Anteroposterior orientation of raw images

Prior to quantification, all images were corrected for the anteroposterior body axis. This axis was determined from brightfield images acquired in parallel by manually annotating the fish midline, and the resulting angle was used to rotate confocal images and downstream measurements so that all samples were aligned to a common anteroposterior reference frame. This registration step was applied uniformly before all subsequent analyses described below, including quantification of sfGFP-Vangl2 distribution and hair-bundle orientations.

#### Shape-normalized sfGFP-Vangl2 intensity maps

Raw images were oriented to a common anteroposterior reference frame as described above and analyzed in Fiji (ImageJ) (33) to quantify the angular distribution of sfGFP-Vangl2 at apical hair-cell boundaries. Hair cells were identified using the HC *βactin* (*Tg:myo6b:actb1-mScarletI*) reporter, and a 10-pixel-thick line ROI was manually traced along each apical cell boundary.

To pool the perimeter distribution of sfGFP-Vangl2 across hair cells of disparate size and shape, each cell’s apical contour was reduced to a single common coordinate frame. The contour was fitted with an ellipse by the direct least-squares method of Halíř and Flusser(34), which returns the best-fitting conic under the constraint 4*ac* − *b*^2^ = 1. The center, the orientation *φ*, and the semi-axes a and b were taken from the fitted conic and from its quadratic form. The cell and the surrounding fluorescence were then rescaled until this ellipse was a circle of unit radius, so that *r* = 1 is the hair-cell boundary, *r* < 1 the cell interior and *r* > 1 its surroundings. For the rare near-degenerate contour, a principal-component fit of the vertices was used as a fallback.

Intensity was then read along 360 rays, one per degree, each running outward from the cell centroid in 1-px steps by bilinear interpolation: through the cell interior, across the traced boundary, and on into a margin beyond it. The margin was round(0.5 × mean(*a, b*)) px wide, roughly half a cell radius, so every ray records the cell, its membrane and the tissue immediately outside. These margins overlap neighboring cells, so each pixel was assigned to whichever cell was nearest (a Voronoi partition) and signal falling in a neighbor’s territory was discarded. Because the margin is set in pixels before rescaling, and rescaling stretches the two axes unequally, its width in normalized units is not constant: rays end at a median radius of 1.60 rather than exactly 1.5 (5th–95th percentile, 1.52–1.72). Only this outer fringe is affected, well beyond the *r* = 1.45 disc that the figures show.

The background-subtracted signal was then sampled by bilinear interpolation along 360 radial rays cast at one-degree spacing from the geometric centroid, each ray running from the cell centroid in 1-px steps out to the traced boundary and a further round(0.5 ×mean(*a, b*)) px beyond it (floor 5 px), roughly half a cell radius, so every ray records the cell, its membrane and the tissue immediately outside. The rescaling map *T* = *R*(+*φ*) · diag(1*/a*, 1*/b*) · *R*(−*φ*) was applied about the fitted ellipse center. This map results in a pure anisotropic stretch that maps the fitted ellipse exactly onto the unit circle and applies no rotation to the image frame so that the anteroposterior axis, made horizontal by the registration described above, carries over unchanged. Within each neuromast, the retained intensities were divided by a shared area-weighted mean before pooling. This normalizing mean was taken over the interior samples of all cells in the neuromast, with each sample weighted by its radial distance from the centroid to ensure that denser ray sampling near the center did not bias the result. Maps were then built in two steps, the neuromast rather than the cell being the unit of replication. First, all points of one neuromast were binned on a 0.05-unit grid, the same grid for every neuromast and condition, each bin holding the mean of its points, and smoothed with a coverage-weighted Gaussian of *σ* = 2 bins (0.10 units); signal and coverage were smoothed separately and divided, so empty bins would not dilute the signal. Second, the neuromast maps of a condition were averaged bin by bin with equal weight. The color scale spans the 2nd to 98th percentiles of all six condition maps together, so the panels are comparable. A dashed circle at unit radius marks the mean fitted cell boundary; the displayed map is a dimensionless per-neuromast relative density. Because each ray is sampled along its whole length, from the center through the membrane and into the halo, rather than only within a narrow band at the traced outline, the maps and profiles are robust to the precision of the manual segmentation. Jitter and small irregularities in the traced outline average out over the 360 rays; the membrane enrichment is recovered at its true radial position, whether the outline falls exactly on the membrane or not; and the anteroposterior asymmetry is essentially independent of how the boundary is drawn. The outline affects only the fitted ellipse that sets the radial scale (*r* = 1).

#### Radial sfGFP-Vangl2 intensity profiles

For each neuromast, an azimuthally averaged radial profile was computed from the same unit-disk-normalized, per-neuromast-normalized point clouds used for the maps. The normalized intensities were binned by radial distance *r* from the cell center in 0.05-unit steps, from the center outward through the *r* = 1 boundary and into the margin beyond the cell, and averaged within each annulus over all angles and all cells of the neuromast. Per-neuromast profiles were then averaged across the neuromasts of a condition, each neuromast weighted equally, and are plotted as the mean ± SEM, with the SEM band shown only where at least two neuromasts contributed. Because the profiles average over angle, they report the radial distribution of sfGFP-Vangl2, that is, the enrichment of Vangl2 at the hair-cell boundary relative to the cell interior and the radial position of that enrichment, but not planar polarity. These profiles are shown in Figure S1C, which overlay control, hair-cell-degradation (HC zGrad), and support-cell-degradation (SC zGrad) neuromasts for *vangl2*^sfGFP/+^ and *vangl2*^sfGFP/sfGFP^ larvae, corresponding to the samples analyzed in Figures 1 and 2, respectively.

#### Anteroposterior sfGFP-Vangl2 enrichment (E_AP_)

To quantify the anteroposterior (AP) asymmetry of sfGFP-Vangl2 localization at the hair-cell membrane, we measured radial intensity profiles along manually traced 10-pixel-thick cell-outline ROIs (Fiji, Multi Plot) in their native, unwarped image coordinates. For each sampled point along a trace, the true planar position on the ROI outline was recovered from the corresponding ROI file, and its polar angle relative to the cell centroid was calculated by arctan 2(Δ*y*, Δ*x*), corrected for the tracing direction used during manual outlining. Per-cell angular intensity profiles were interpolated onto a common 360°grid, averaged (un-weighted) across all hair cells within a neuromast, and the resulting neuromast-level profile was normalized to its own total intensity after background subtraction. Anterior and posterior signals were defined as the sum of the normalized intensities within symmetric *±*45°sectors centered on the anterior (0°) and posterior (180°) poles of the AP axis, respectively. The AP enrichment index was calculated per neuromast as

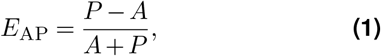

where *A* and *P* represent the summed normalized intensity within anterior and posterior sectors. *E*_AP_ is bounded within [−1, +1] by construction; positive values indicate posterior enrichment, and negative values indicate anterior enrichment in Figures 1 and 2. Differences between conditions were assessed using Kruskal–Wallis tests followed by pairwise Mann–Whitney tests with Holm correction for multiple comparisons.

#### Hair-bundle orientation measurement and analysis

Hair-bundle polarity was quantified by annotating a line from the distal edge of each hair bundle toward the kinocilium and measuring its orientation relative to the anteroposterior axis (see Figure 1E). Young hair cells with ambiguous bundle morphology or orientation were excluded from analysis.

Rose plots were generated using 15°bins with wedge areas proportional to the fraction of bundles per bin. Axis limits were held constant within each comparison. Because hair-bundle polarity is axial (i.e., orientations separated by 180°represent the same polarity axis), analyses were performed using doubled angles. Alignment of bundle orientations along the AP axis was quantified using the axial resultant length, which forms the basis for axial Rayleigh tests(35),

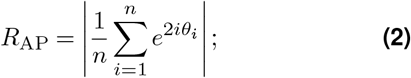

where *θ*_*i*_ represents the orientation angle of the *i*-th hair bundle and *n* is the number of observations. *R*_AP_ ranges from 0 (random orientation) to 1 (perfect alignment along the AP axis). *R*_AP_ values were calculated per neuromast. Differences in alignment strength between conditions were assessed using Kruskal–Wallis tests followed by pairwise Mann–Whitney tests with Holm correction for multiple comparisons (Figures 1, 2, and 4) or paired Wilcoxon signedrank tests where indicated (Figure 3). Departure from uniformity of orientation distributions was evaluated using axial Rayleigh tests applied to doubled angles.

## Acknowledgments

We dedicate this paper to the memory of A.J.H., whose mentorship and curiosity shaped this work and continue to inspire us. We thank members of the Hudspeth laboratory for valuable advice and suggestions on the manuscript. Amy Shyer, Gregory Alushin, Michael Harrison, and Tim Stearns provided scientific advice and helpful discussions throughout the development of the project. We are grateful to members of the Bio-Imaging Resource Center at The Rockefeller University for their advice on image acquisition and analysis. We greatly appreciate Samantha Campbell’s expert care of the fish facility. S.J. was supported by a Medical Scientist Training Program grant from the National Institute of General Medical Sciences of the National Institutes of Health under award number T32GM152349 to the Weill Cornell/Rockefeller/Sloan Kettering Tri-Institutional MD-PhD Program. F.G. was supported by a Kavli Neural Systems Institute Fellowship. We thank the Howard Hughes Medical Institute for supporting this work. A.J. is grateful to Biohub donors P. Chan and M. Zuckerberg for their generous support.

## Author contributions

S.J. and A.J.H. conceptualized the project, and A.J.H., A.J., and A.S. provided ongoing scientific feedback. M.J. generated the vangl2^sfGFP/sfGFP^ and *Tg(βactin2:loxP-mCherry-STOP-loxP-zGrad)* zebrafish lines in B.C.’s laboratory and provided these lines together with *Tg(hsp:Cre)*. S.J. generated the *Tg(myo6b:actb1-mScarletI), Tg(myo6b:zGrad-2A-mCherry)*, and *Tg(she:zGrad-2A-mCherry)* transgenic lines used in this study. S.J. performed the experiments and analyzed the data. F.G. developed the methodology and code for Vangl2 radial intensity profiling and generated the resulting plots; S.J. generated all other plots and assembled the figures. S.J. wrote the original manuscript with input from A.S. and A.J. All authors except A.J.H. reviewed and edited the manuscript. The project was supervised by A.J.H., F.G., A.S., and A.J. A.J.H. and B.C. acquired funding and provided resources.

## Resource availability

### Lead contact

Requests for further information and resources should be directed to and will be fulfilled by the lead contact, Adrian Jacobo.

### Materials availability

All reagents and fish lines generated in this study are available from the lead contact without restriction.

### Data and code availability

The data and code generated for this study are available from the lead contact without restriction.

## Declaration of generative AI and AI-assisted technologies in the writing process

During the preparation of this work, the authors used Chat-GPT (OpenAI) to revise grammar and sentence structure. After using this tool, the authors reviewed and edited the content as needed and take full responsibility for the content of the publication.

## Supplemental information

Document S1. Supplemental Figures S1–S3.

