## Supplementary Information for "Vangl2 acts in distinct cell types to establish bidirectional hair-bundle polarity and maintains tissue-wide alignment in zebrafish neuromasts"

### Document S1. Supplemental Figures S1–S3

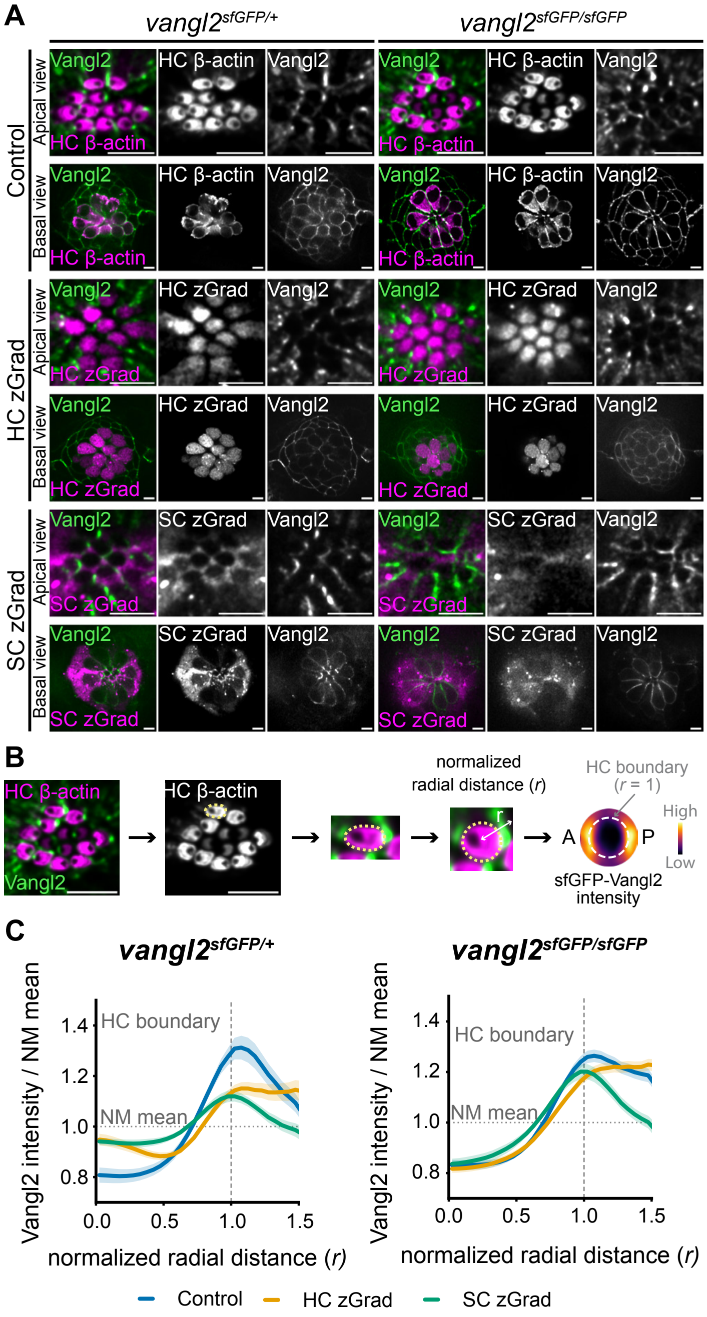


**Figure S1. Characterization of cell-type-specific zGrad expression and radial Vangl2 distribution relative to hair-cell boundaries.** Related to Figures 1 and 2. (A) Representative apical and basal images of 5 dpf *vangl2*$^{sfGFP/+}$ or *vangl2*$^{sfGFP/sfGFP}$ neuromasts under control conditions or after zGrad-mediated degradation of sfGFP-Vangl2 in hair cells (HCs; HC zGrad, *Tg(myo6b:zGrad-2A-mCherry)*) or support cells (SCs; SC zGrad, *Tg(she:zGrad-2A-mCherry)*), as indicated. In control neuromasts, HC $\beta$-actin from *Tg(myo6b:actb1-mScarletI)* marks the stereotyped organization of HCs at apical and basal planes: apically, HCs are tightly packed, whereas basally, HC bodies form a central cluster. In HC zGrad neuromasts, zGrad-2A-mCherry follows the HC $\beta$-actin pattern seen in controls, marking apical HC positions and the basal HC cluster, with reduced sfGFP-Vangl2 signal in this region. In SC zGrad neuromasts, zGrad-2A-mCherry shows a complementary pattern, surrounding apical HC positions and interspersing around the HC region in basal planes, with sfGFP-Vangl2 signal remaining more prominent in the basal HC cluster. Together, these complementary zGrad-2A-mCherry and sfGFP-Vangl2 patterns support the interpretation of cell-type-enriched zGrad activity and residual Vangl2 distribution. (B) Workflow for generating shape-normalized sfGFP-Vangl2 maps. Because HCs and SCs are tightly interdigitated at the apical surface, sfGFP-Vangl2 signal was quantified relative to HC apical boundaries. HC boundaries were manually segmented using HC $\beta$-actin from *Tg(myo6b:actb1-mScarletI)* and mapped onto a common unit-circle coordinate frame by an anisotropic stretch along each cell's principal axes; the anteroposterior axis is that of the image, which was pre-registered to the fish's anteroposterior body axis. Background-subtracted sfGFP-Vangl2 intensity was then sampled over both angle and normalized radial distance ($r$) and pooled by condition after neuromast-level normalization. Color shows the spatial distribution of Vangl2 enrichment (brighter color means higher enrichment) relative to the neuromasts' average; all panels share one scale. The normalized HC boundary corresponds to $r$ = 1. (C) Azimuthally averaged radial profiles of sfGFP-Vangl2 intensity relative to the normalized HC boundary in *vangl2*$^{sfGFP/+}$ and *vangl2*$^{sfGFP/sfGFP}$ neuromasts. Because these profiles average over angle, they report Vangl2 enrichment at the membrane relative to the cell interior and exterior, that is, the position and sharpness of the boundary ring. Profiles show neuromast-normalized sfGFP-Vangl2 intensity as a function of normalized radial distance (r), with $r$ = 1 marking the HC boundary. Horizontal dashed lines indicate the neuromast mean. In SC zGrad neuromasts, residual sfGFP-Vangl2 remains enriched near the HC boundary, consistent with retained HC-associated Vangl2. In HC zGrad neuromasts, residual sfGFP-Vangl2 is shifted farther from the HC boundary, consistent with remaining SC-associated Vangl2. This radial separation of the two residual pools indicates that the two degrons deplete Vangl2 from different cellular sources. For C, *vangl2*$^{sfGFP/+}$ and *vangl2*$^{sfGFP/sfGFP}$ samples are the same as those analyzed in Figures 1 and 2, respectively. Scale bars, 5 µm.

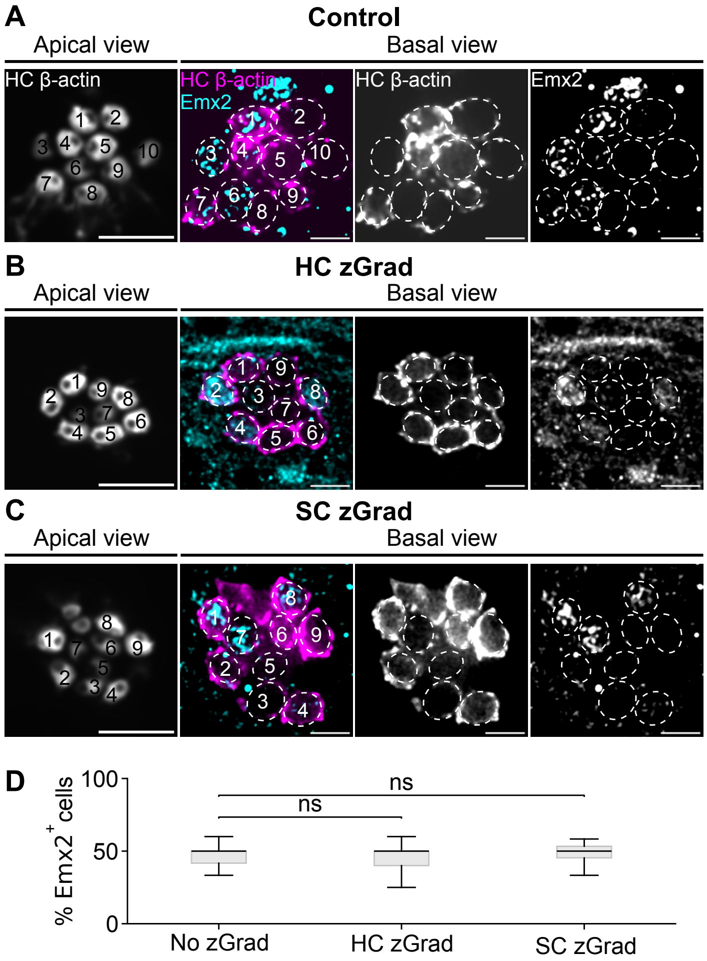


**Figure S2. Hair-cell identity proportions are unchanged after cell-type-specific Vangl2 degradation.** Related to Figure 3. (A–C) Representative 5 dpf *vangl2*$^{sfGFP/sfGFP}$ neuromasts under control conditions (A, no zGrad), after zGrad-mediated degradation of sfGFP-Vangl2 in hair cells (HCs; B, HC zGrad, *Tg(myo6b:zGrad-2A-mCherry)*), or after degradation in support cells (SCs; C, SC zGrad, *Tg(she:zGrad-2A-mCherry)*). Apical views show HC $\beta$-actin used to identify and number HCs; basal views show HC $\beta$-actin and nuclear Emx2 signal used to assign Emx2$^+$ and Emx2$^-$ HC identities. Numbered HCs were matched across apical and basal z-planes by scrolling through the full z-stack. (D) Percentage of Emx2$^+$ HCs per neuromast across conditions. For D, n = 315 HCs/33 neuromasts/16 fish for control, 555 HCs/58 neuromasts/32 fish for HC zGrad, and 282 HCs/23 neuromasts/13 fish for SC zGrad. Boxes show median and interquartile range. Statistical significance was assessed using two-sided Mann–Whitney U tests on neuromast-level percentages. Exact p-values: control vs HC zGrad, P = 0.740; control vs SC zGrad, P = 0.552; HC zGrad vs SC zGrad, P = 0.288. ns, not significant. Scale bars, 5 µm.

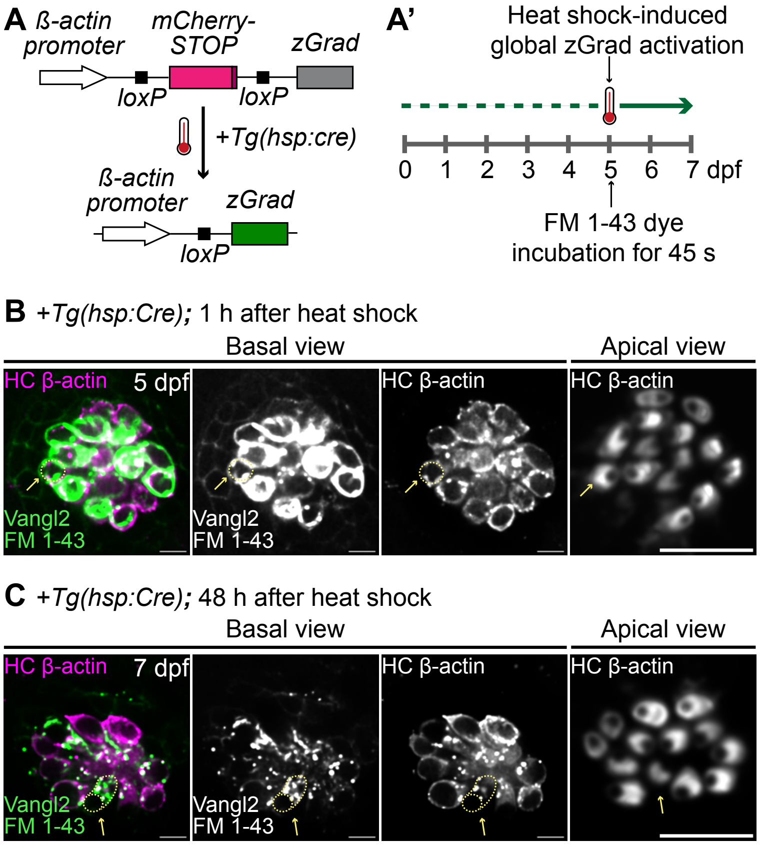


**Figure S3. FM1-43 styryl dye retention provides evidence for hair-bundle misorientation in pre-existing hair cells after late Vangl2 degradation.** Related to Figure 4. (A) Schematic of heat-shock-induced global zGrad activation in *Tg(βactin2:loxP-mCherry-STOP-loxP-zGrad); Tg(hsp:Cre)* larvae. Heat-shock-induced Cre excises the mCherry-STOP cassette, placing zGrad under the control of the $\beta$-actin promoter. (A′) Experimental timeline. Larvae were heat shocked at 5 dpf to induce global zGrad expression and treated immediately and transiently with FM1-43 to label hair cells (HCs) present at the time of perturbation. Larvae were imaged either 1 h or 48 h after heat shock. (B, C) Representative basal and apical images 1 h (B) or 48 h (C) after late heat shock. HC $\beta$-actin from *Tg(myo6b:actb1-mScarletI)* marks HCs and hair bundles (HBs). sfGFP-Vangl2 and FM1-43 were detected in the same green channel; sfGFP-Vangl2 signal was still detectable 1 h after heat shock and largely depleted by 48 h. At 48 h after heat shock, the retained FM1-43 signal appeared punctate but remained associated with HC $\beta$-actin-positive cells in basal planes, while apical HC $\beta$-actin images showed misaligned HBs. Yellow dashes and arrows identify the same HC with retained FM1-43 in the basal view and apical HB-orientation in the corresponding apical view in each panel, respectively. Together, FM1-43 retention and HC $\beta$-actin labeling support that HCs present at the time of late heat shock persisted and displayed hair-bundle misorientation after Vangl2 degradation. Scale bars, 5 µm.
